# BDNF-TrkB signaling promotes synaptic GluN2A-NMDA receptor expression and network hyperexcitability in cultured hippocampal neurons and during status epilepticus

**DOI:** 10.1101/2025.10.13.682007

**Authors:** Pasqualino De Luca, Miranda Mele, Francesca Napoli, Philemon Mshelia, Rui O. Costa, Carlos B. Duarte

## Abstract

Brain-derived neurotrophic factor (BDNF) is a key modulator of synaptic function, acting through activation of TrkB receptors. This neurotrophic factor mediates synaptic plasticity and plays an important role in epileptogenesis, but the underlying molecular mechanisms have not been fully elucidated. In this work, we investigated the role of BDNF-TrkB signaling in the regulation of synaptic GluN2A-containing NMDA receptors (NMDAR), and the impact on network synchronization in cultured hippocampal neurons. Incubation with BDNF increased the synaptic surface expression of GluN2A-containing NMDAR in rat hippocampal synaptoneurosomes and in cultured hippocampal neurons. The effect in the latter preparation was time-dependent and required new protein synthesis. Mechanistically, we identified a signaling cascade involving hnRNPK, the non-receptor tyrosine kinase Pyk2, and protein kinase C (PKC) as essential for mediating BDNF-induced upregulation in the synaptic expression of GluN2A-containing NMDAR. Knockdown of hnRNPK or Pyk2, pharmacological inhibition of PKC, or expression of a phosphorylation -deficient Pyk2 mutant abolished BDNF-induced synaptic surface accumulation of Glu2A. Moreover, Pyk2 phosphorylation at Y402 was necessary for both basal and BDNF-induced synaptic GluN2A expression. Functional multielectrode array (MEA) recordings showed that endogenous BDNF and GluN2A-containing NMDAR contributed to the increase in network activity in cultured hippocampal neurons evoked by stimulation with a cocktail including bicuculline, 4-aminopyridine, and glycine. Importantly, BDNF-TrkB signaling also mediated the upregulation in the hippocampal synaptic surface expression of GluN2A-containing NMDAR, as determined in rats subjected to the pilocarpine model of temporal lobe epilepsy, where increased GluN2A synaptic expression was observed and shown to be TrkB-dependent. These findings unveil a crucial BDNF/TrkB-PKC-Pyk2-hnRNPK signaling axis that regulates synaptic GluN2A levels and network excitability, offering novel insights into the molecular basis of synaptic plasticity.

## Introduction

Brain-derived neurotrophic factor (BDNF) is a critical mediator in neuronal development, synaptic plasticity, and cognitive function (Carvalho, Caldeira et al. 2008, Park and Poo 2013, Leal, Comprido et al. 2014, Kowianski, Lietzau et al. 2018, Mohandas, Rishikesh et al. 2025). Through activation of its high-affinity receptor TrkB, BDNF mediates multiple aspects of excitatory synaptic transmission, including long-term synaptic potentiation (LTP) (Korte, Carroll et al. 1995, Carvalho, Caldeira et al. 2008, Leal, Comprido et al. 2014, Mohandas, Rishikesh et al. 2025) and epileptogenesis (Gu, Huang et al. 2015, Huang, He et al. 2019, Krishnamurthy, Huang et al. 2019). Stimulation of TrkB receptors leads to downstream activation of the Ras/ Extracellular signal-regulated kinases (ERK) and phosphatidylinositol 3-kinases (PI3-K)/Akt signaling pathways, together with phospholipase C-γ (PLC-γ). Studies performed in mice expressing targeted mutations in the PLC-γ binding sites of TrkB showed a role for this pathway in LTP of CA1 synapses and in associative learning (Minichiello, Calella et al. 2002, Gruart, Sciarretta et al. 2007), and these effects are mediated at pre- and post-synaptic level (Gartner, Polnau et al. 2006). Interestingly, transient inhibition of TrkB-PLCγ signalling using a specific peptide prevented epilepsy induced by status epilepticus in mice (Gu, Huang et al. 2015), showing a key role for this pathway in BDNF-induced potentiation of glutamatergic synapses. Stimulation of local translation is also an important feature in the potentiation of CA1 synapses by BDNF (Kang and Schuman 1996, Schratt, Nigh et al. 2004, Takei, Inamura et al. 2004, Leal, Comprido et al. 2014, Costa, Martins et al. 2022), and it is also required for consolidation of LTP in the dentate gyrus following infusion of the neurotrophin (Messaoudi, Kanhema et al. 2007, Panja, Kenney et al. 2014).

N-methyl-D-aspartate receptors play important roles in the plasticity of glutamatergic synapses. These are heterotetrameric receptors containing in most cases, two obligatory GluN1 subunits, possessing the glycine/D-serine binding sites, and two GluN2 subunits which bind glutamate. Among the NMDAR subunits, GluN2A and GluN2B are predominantly expressed in the adult hippocampus (Shipton and Paulsen 2014), possessing distinct biophysical properties, including the affinity for the agonist, open probability, Ca^2+^ permeability and charge transfer, and deactivation kinetics (Yashiro and Philpot 2008). Given the differences between the C-terminus of GluN2A and GluN2B, the two subunits are also differentially regulated (Tahiri, Corti et al. 2025). This is further suggested by the largely non-overlapping nanoscale distribution of the two NMDAR subunits in hippocampal synapses (Zeng, Shang et al. 2016, Kellermayer, Ferreira et al. 2018). Together, these differences contribute to a specific coupling to the regulation of downstream signaling pathways that influence neuronal function (Paoletti, Bellone et al. 2013, Tian, Stroebel et al. 2021, Ladagu, Olopade et al. 2023, Storey, Riquelme et al. 2025). GluN2B is predominantly expressed during early postnatal development and is often found at extrasynaptic sites, while GluN2A expression increases during maturation and is enriched at synaptic locations (Liu, Wong et al. 2004, Yashiro and Philpot 2008). The developmental switch from GluN2B to GluN2A subunits is a key event in the functional maturation of glutamatergic synapses and is thought to influence the direction and magnitude of long-term synaptic changes, including LTP, which underlie certain forms of learning and memory formation (Paoletti, Bellone et al. 2013, Franchini, Carrano et al. 2020). A rapid surface accumulation of NMDAR was also observed in dentate gyrus granule cells, as well as in CA3 pyramidal neurons, after 1h of lithium-pilocarpine status epilepticus in rats (Naylor, Liu et al. 2013). In addition, an upregulation of GluN2B-containing NMDAR (GluN2B-NMDAR) was observed in the lateral perforant path synapses with hippocampal dentate granule neurons, in studies performed using the pilocarpine model of temporal lobe epilepsy (Klatte, Kirschstein et al. 2013). An increase in surface GluN2B was also observed in hippocampal synaptoneurosomes under similar conditions, by a mechanism sensitive to the TrkB receptor inhibitor ANA-12 (De Luca, Mele et al. 2025).

BDNF upregulates the activity of postsynaptic NMDAR in cortical and hippocampal CA1 pyramidal neurons (Kolb, Trettel et al. 2005), in dentate gyrus granule cells (Madara and Levine 2008), as well as in cultured hippocampal neurons (Leal, Comprido et al. 2017, Afonso, De Luca et al. 2019, De Luca, Mele et al. 2025). Previous studies also showed that BDNF modulates the surface expression and synaptic incorporation of GluN2B-NMDAR via mechanisms involving protein synthesis (Xu, Plummer et al. 2006, Caldeira, Melo et al. 2007). Although the role of BDNF in modulating NMDAR function is well established, much less is known about its influence on GluN2A-containing receptors (GluN2A-NMDAR), which are predominantly localized in the synaptic compartment. Understanding how BDNF regulates the synaptic expression of GluN2A -NMDAR should provide important insights into the mechanisms of synaptic modulation by this neurotrophin, including the synaptic alterations associated with distinct forms of long-term potentiation (LTP) and epilepsy. In this study, we demonstrate a critical role for BDNF-TrkB signalling in enhancing the synaptic expression of GluN2A-NMDAR, a process accompanied by increased synchronized activity in cultured hippocampal neurons and by hyperexcitability during status epilepticus. Furthermore, we identified the TrkB-PKC-Pyk2-hnRNPK signalling axis as the key mediator of these effects of BDNF.

## Material and Methods

### Animals

Wistar rats and male Sprague-Dawley rats, aged 6-8 weeks (Charles River Laboratories, Barcelona, Spain), were maintained under controlled temperature (21 ± 1°C) and humidity (55 ± 10%) conditions, with a 12:12 h light/dark cycle and access to food and water ad libitum. Animals were handled according to Portuguese Law (DL 113/2013) and the European Community Guidelines (Directive, 2010/63/EU) on the protection of animals for scientific experimentation.

### Isolation of synaptoneurosomes

Synaptoneurosomes were isolated as previously described (De Luca, Mele et al. 2025). Briefly, two hippocampi were dissected from adult Sprague-Dawley rats (6-8 weeks old). The tissue was minced with a blade and homogenized with Kontes Dounce Tissue Grinder in a buffer containing 0.32 M sucrose, 10 mM HEPES-Tris pH 7.4, and 0.1 mM EGTA pH 8, using first a pestle of 0.889-0.165 mm clearance, followed by a pestle of 0.025-0.076 mm. After centrifugation for 3 min at 1000 x g at 4°C, the collected supernatant was passed through triple layered nylon membranes (150 and 50 μm, VWR) and then through an 8 μm pore size filter (Millipore, MA). The flow-through was centrifuged for 15 min at 10000 x g, and the resulting pellet was suspended in the buffer used for homogenization. All the procedure was performed on ice or at 4 °C. Part of the synaptoneurosome preparation was plated on glass coverslips and processed for immunostaining, and the remaining material was used for protein extraction followed by western blot analyses (see below).

### Hippocampal cultures

Neuronal low-density hippocampal cultures were prepared from the hippocampi of E18-E19 Wistar rat embryos, after incubation with trypsin (0.06% (w/v), GIBCO - Life Technologies) in Ca^2+^- and Mg^2+^-free Hanks’ balanced salt solution (HBSS: 5.36 mМ KCl, 0.44 mM KH_2_PO_4_, 137 mM NaCl, 4.16 mM NaHCO_3_, 0.34 mM Na_2_HPO_4_·2H_2_O, 5 mM glucose, 1 mM sodium pyruvate, 10 mM HEPES, and 0.001% phenol red), for 15 min at 37°C (Afonso, De Luca et al. 2019). The hippocampi were then washed with Hanks’ balanced salt solution containing 10% fetal bovine serum (GIBCO - Life Technologies) to stop trypsin activity and transferred to Neurobasal medium (GIBCO - Life Technologies) supplemented with SM1 supplement (1:50 dilution, STEMCELL Technologies), 25 μM glutamate, 0.5 mM glutamine, and 0.12 mg/ml gentamycin (GIBCO - Life Technologies). Neurons were plated at a final density of 1.4 × 10^4^ cells/cm^2^ on poly-D-lysine (PDL)-coated glass coverslips in neuronal plating medium (Minimum Essential Medium [MEM, Sigma-Aldrich] supplemented with 10% horse serum, 0.6% glucose and 1 mM pyruvic acid). After 2 h the coverslips were flipped over an astroglia feeder layer in Neurobasal medium (GIBCO - Life Technologies) supplemented with SM1 supplement (1:50 dilution, STEMCELL Technologies), 25 μM glutamate, 0.5 mM glutamine and 0.12 mg/ml gentamycin (GIBCO - Life Technologies). The neurons grew face down over the feeder layer but were kept separated from the glia by paraffin dots on the neuronal side of the coverslips. To prevent the overgrowth of glial cells, neuronal cultures were treated with 10 μM 5-Fluoro-2′-deoxyuridine (FDU) (Sigma-Aldrich) at DIV 3. Cultures were maintained in a humidified incubator with 5% CO_2_/95% air at 37°C for up to 2 weeks, feeding the cells once per week with the same Neurobasal medium described above, but without added glutamate. At DIV 14-15 the neurons were stimulated with 50 ng/ml BDNF (Peprotech) for the indicated period. Where indicated, cells were pre-treated for 45 min with a protein synthesis inhibitor, 40-45 min with a protein kinase C inhibitor, (3-[1-[3-(Dimethylamino)-propyl]-5-methoxy-1H-indol-3-yl]-4-(1H-indol-3-yl)-1H-pyrrole-2,5-dione (GӦ6983, 100 nM) or with the vehicle dimethyl sulfoxide (DMSO, 1:1000 dilution, Sigma-Aldrich), as control.

### Staining of surface GluN2A-NMDAR in cultured hippocampal neurons and quantitative image analysis

To label surface GluN2A-NMDAR, live neurons (low-density hippocampal cultures) were incubated for 10 min at room temperature with an antibody against an extracellular epitope of the GluN2A N-terminus (1:100; AGC-002, Alomone Labs) diluted in a saline buffer (145 mM NaCl, 5 mM glucose, 10 mM HEPES, 5 mM KCl, 1.8 mM CaCl_2_, 1 mM MgCl_2,_ [pH 7.3]), as previously described (Afonso, De Luca et al. 2019). Neurons were then fixed for 15 min in 4% sucrose and 4% paraformaldehyde in phosphate-buffered saline (PBS: 137 mM NaCl, 2.7 mM KCl, 1.8 mM KH2PO_4_, 10 mM Na_2_HPO_4_·2H_2_O [pH 7.4]) at room temperature and permeabilized with PBS + 0.3% (v/v) Triton X-100 for 5 min at 4°C. The preparations were then incubated in 10% (w/v) bovine serum albumin (BSA) in PBS for 1 h at room temperature to block nonspecific staining and incubated with the appropriate primary antibody (anti-MAP2 [1:10,000; ab5392, Abcam], anti–PSD-95 [1:200; 7E3-1B8, Thermo Scientific], and anti-vGluT1 [1:5000; AB5905, Millipore]) diluted in 3% (w/v) BSA in PBS (overnight, 4°C). After washing six times in PBS, cells were incubated with the appropriate secondary antibody (Alexa Fluor 488-conjugated anti-mouse [1:1000; A-11001, Thermo Fisher Scientific], Alexa Fluor 488-conjugated anti-rabbit [1:1000; A-11034, Thermo Fisher Scientific], Alexa Fluor 568-conjugated anti-mouse [1:1000; A-11004, Thermo Fisher Scientific], Alexa Fluor 568-conjugated anti-rabbit [1:1000; A-11036, Thermo Fisher Scientific], Alexa Fluor 647-conjugated anti-guinea pig [1:500; A-21450, Thermo Fisher Scientific], and AMCA (aminomethyl-coumarin)-conjugated anti-chicken [1:200; #103-155-155, Jackson ImmunoResearch]) diluted in 3% (w/v) BSA in PBS (1 h at room temperature). The coverslips were mounted using a fluorescent mounting medium (DAKO).

Fluorescence imaging was performed on a Zeiss Axio Observer Z.1 microscope using a 63 × 1.4 numerical aperture (NA) oil objective and a Zeiss Axio Imager Z.2 microscope using a 63× 1.4 NA oil objective, both equipped with a Zeiss HRm AxioCam. Images were quantified using the Fiji image analysis software. For quantification, sets of cells were cultured and stained simultaneously and imaged using identical settings. The region of interest was randomly selected, and the dendritic length was measured based on MAP2 staining. The protein signals were analyzed after setting the appropriate thresholds, and the recognizable puncta under those conditions were included in the analysis. For each experiment, similar threshold levels were used to quantify the number and the integrated intensity of puncta in dendrites. Measurements were performed in three to six independent preparations, as indicated in the figure captions.

To analyze GluN2A synaptic surface expression, the PSD-95 and vGluT1 signals were thresholded and their colocalization was determined. The surface GluN2A signal was measured after thresholds were set so that recognizable puncta were included in the analysis. The surface GluN2A signal present in glutamatergic synapses was obtained by measuring the surface GluN2A puncta positive for both PSD-95 and vGluT1. The results were represented *per* density of excitatory synapses (number of positive PSD-95-vGluT1 puncta that colocalized per dendritic length). Fluorescence imaging was performed at the MICC Imaging facility of CNC-UC, partially supported by PPBI – Portuguese Platform of BioImaging (PPBI-POCI-01-0145-FEDER-022122).

### Live immunostaining of surface GluN2A-NMDAR in hippocampal synaptoneurosomes

Synaptoneurosomes (35 µl) were plated on 10 mm diameter coverslips coated with poly-D-lysine (0.1 mg/mL) and left in a humidified chamber for 1 h at room temperature, to allow adhesion. Where indicated, synaptoneurosomes were incubated with BDNF (50 ng/mL, Peprotech) at 30°C during this period, and control experiments in the absence of the neurotrophin were also performed. Immediately after stimulation, synaptoneurosomes were processed for GluN2A live immunostaining as described below (see also (Masella, Silva et al. 2024, De Luca, Mele et al. 2025)). Live staining was performed as described above for the analysis of surface GluN2A expression in cultured hippocampal neurons (Afonso, De Luca et al. 2019, Masella, Silva et al. 2024).

### Fluorescence imaging acquisition and quantitative analysis of NMDAR surface staining in synaptoneurosomes

Fluorescence imaging of synaptoneurosomes was conducted using a Carl Zeiss Axio Imager Z2 widefield fluorescence microscope, as previously described (Masella, Silva et al. 2024, De Luca, Mele et al. 2025). Images were captured with a Plan-Apochromat 100×/1.4 numerical aperture oil objective and a Zeiss HRm AxioCam camera. Zeiss filter sets 31, 38 (HE), and 50 were used, and phase-contrast images were also obtained. Consistent exposure times and light intensities were maintained across different experimental conditions. Image analysis was performed with ImageJ software (National Institutes of Health, MD, USA) using a custom-designed macro for this experiment. Intact synaptoneurosomes were identified based on the following criteria: juxtaposition of presynaptic (VGluT1) and postsynaptic (PSD-95) protein clusters, visibility as dark objects in phase-contrast images indicating sealed synaptoneurosomes, size range (300–1000 nm), and a snowman-like shape as previously described (Tai, Wang et al. 2014). Approximately 500–600 synaptoneurosomes per condition were manually delineated as regions of interest (ROIs), and background and thresholds were set for GluN2A signals. The analysis quantified the area mean intensity as well as the integrated density of GluN2A staining within the synaptoneurosomes.

### Lithium-pilocarpine model of Status Epilepticus

The protocol to induce status epilepticus using the lithium-pilocarpine model was approved by DGAV - Direcção Geral de Ambiente e Veterinária (Ref. 0421/000/000/2020). Experiments were conducted using 6-8 weeks-old male Sprague-Dawley rats obtained from Charles River Laboratories (Barcelona, Spain) as previously described (De Luca, Mele et al. 2025). First, the animals were weighed to determine the correct dosage of each substance. Lithium chloride (LiCl; 127 mg/ml in deionized sterile water) was administered first at a dosage of 127 mg/kg via intraperitoneal injection. Twenty hours later, methyl-scopolamine (2 mg/mL in saline [0.9% NaCl]) was administered at a dosage of 1 mg/kg, also via intraperitoneal injection. The animals were maintained for 45 min more before intraperitoneal injection of pilocarpine (20 mg/mL in saline [0.9% NaCl]) at a dosage of 10 mg/kg. During this period, the animals were closely monitored for any adverse reactions. Behavioural changes and physiological responses were recorded with a video camera, with the severity of seizures classified using the Racine scale (Racine 1972) that categorizes the progression of seizure severity based on observable behaviours. If an animal did not reach Racine stage 5, indicative of status epilepticus (SE), within 30 min of the pilocarpine injection, an additional dose of pilocarpine (10 mg/kg) was administered. This procedure was repeated every 30 min, up to a maximum of five times, until SE was achieved. A saline control group followed the same protocol as the pilocarpine-injected group, but with saline injections replacing pilocarpine administration at equivalent volumes.

For experiments performed in the presence of the compound ANA-12, the drug was injected (i.p.) at a concentration of 0.5 mg/mL (prepared in saline, 5% DMSO) one hour before the first pilocarpine injection. ANA-12 was administered at a dosage of 0.5 mg/kg. Following this procedure, the animals were maintained for 15 min before administration of methyl-scopolamine, and pilocarpine was administered 45 min later. A vehicle control group received an equivalent volume of a saline/DMSO solution, administered intraperitoneally, and based on the volume used for the injection of ANA-12. Rats were anesthetized with isoflurane and euthanized by decapitation, 90 min after the onset of SE determined by the rat’s entry into Stage 5 of seizures according to the Racine scale (Racine 1972). The hippocampi were then dissected and processed for synaptoneurosome isolation as described above.

### Microelectrode Arrays analyses

#### Recording

The cultures for microelectrode array experiments were established on CytoView MEA 24 plates (Axion Biosystems), containing 16 electrodes, with a diameter of 50 μm, spacing of 350 μm and a recording area of 1.1 mm x 1.1 mm, arranged in a 4×4 grid on the bottom of each well. Primary hippocampal rat neurons were plated at a density of 22 × 10^3^ cells/mm^2^, placed directly over the electrode field within each well, precoated with PDL, diluted in 10 μL of Neurobasal medium and laminin (Sigma, 10 μg/mL). Cells were left for 2h at 37°C in a humidified environment to allow adequate attachment, then 500 µL of Neurobasal medium supplemented with SM1 supplement (1:50), 25 μM glutamate, 0.5 mM glutamine and 0.12 mg/ml gentamycin were slowly added to each well in a 2-step process to avoid detaching the cells. Three days later, the culture medium was supplemented with 10 μM FDU. After 7 DIV, 1/4 of the media was replaced with fresh BrainPhys™ Neuronal Medium (Stem Cell Technologies), supplemented with SM1 and gentamicin, twice a week.

Spontaneous network activity from hippocampal cultures grown on MEA plates was recorded using Maestro Edge MEA and Impedance system (Axion Biosystems), coupled to AxIS Navigator software (Axion Biosystems). The raw data were filtered with a Butterworth band-pass filter of 300-5000 Hz, and a threshold for spike detection was set to 5.5 times the standard deviation of the filtered signal. Electrodes were considered active if the mean firing rate during the measurement was at least 5 spikes/min. The basal spontaneous activity of neuronal cells cultured on MEA plates was recorded at 16 DIV (baseline recording, 3 min). After baseline recording, a chemical synaptic stimulation was performed (2.5 mM 4-aminopyridine [4-AP], 50 μM Bicuculline, 10 μM Glycine) and neuronal activity was recorded every 10 min for 30 min. In the experiments performed in the presence of TCN 213 (10 μM) and/or TrkB-FC (1 μg/mL), the inhibitors were added to the medium 30 min before synaptic stimulation, and neuronal activity was recorded 30 minutes later (Baseline 2). Before and during the recordings, the plate was maintained at 37°C and with an atmosphere of 5% CO_2_/95% air. All recordings from the MEA were performed in BrainPhys medium for 3 min.

#### Data analysis

Data obtained from MEA recordings was analyzed using Neural Metrics Tool and AxIS Metric Plotting Tool software (Axion Biosysteams). Raw data files (*.raw) were converted to spike files (*.spk) to allow further processing and analysis of the data. The spike train raster plots were examined with the Neural Metrics Tool to verify the robustness of the activity, as well as the quality and consistency of the well’s activity. Single electrode burst activity was detected using an Inter-Spike Interval (ISI) Threshold set as a minimum of 5 spikes in a maximum 100 ms ISI. Network Burst activity was defined with the Envelope algorithm using the following settings: Threshold Factor = 1.25; Min Inter-Burst Interval (IBI) = 100 ms; Min Electrodes = 70%; Burst Inclusion = 65%.

### Statistical analysis

The results are presented as means ± SEM. Comparisons between multiple groups were performed as indicated in the figure captions, using the Prism 8 software (GraphPad).

## Results

### BDNF increases the synaptic surface expression of GluN2A-NMDAR

Previous studies performed in cultured hippocampal neurons showed that the BDNF-induced upregulation in NMDAR activity is associated with an enhanced synaptic accumulation of GluN2B-NMDAR (Leal, Comprido et al. 2017, Afonso, De Luca et al. 2019, De Luca, Mele et al. 2025), but it remains to be determined whether BDNF also affects the synaptic content in GluN2A under the same conditions. In this work we analysed the effects of BDNF on the surface expression of GluN2A-NMDAR in cultured hippocampal neurons by live-staining with an antibody against an extracellular epitope of GluN2A. After fixation and permeabilization of the plasma membrane, the cells were incubated with antibodies against the dendritic marker microtubule-associated protein 2 (MAP2), and the pre- and post-synaptic markers vGluT1 and PSD-95, respectively. Cultured hippocampal neurons were analyzed for the number (Fig. 1A, B), area (Fig. 1A, C), and intensity (Fig. 1A, D) of GluN2A puncta per dendritic length (as determined based on MAP2 staining). The effect of BDNF on the synaptic surface abundance of GluN2A was assessed by quantifying the number (Fig. 1A, E), area (Fig. 1A, F), and intensity (Fig. 1A, G) of puncta that co-localized with PSD-95 and vGluT1. Neurons incubated with BDNF for 10 min showed a significant upregulation in the total surface expression of GluN2A, as determined by the total number (Fig. 1A, B), area (Fig. 1A, C), and intensity (Fig. 1A, D) of the signal, and similar results were obtained when the synaptic staining was analyzed (Fig. 1A, E-G). This effect was transient since the BDNF-induced increase in GluN2A surface expression was not observed when the cells were incubated with the neurotrophin for 30 min (Fig. 1A-G).

**Figure 1.**
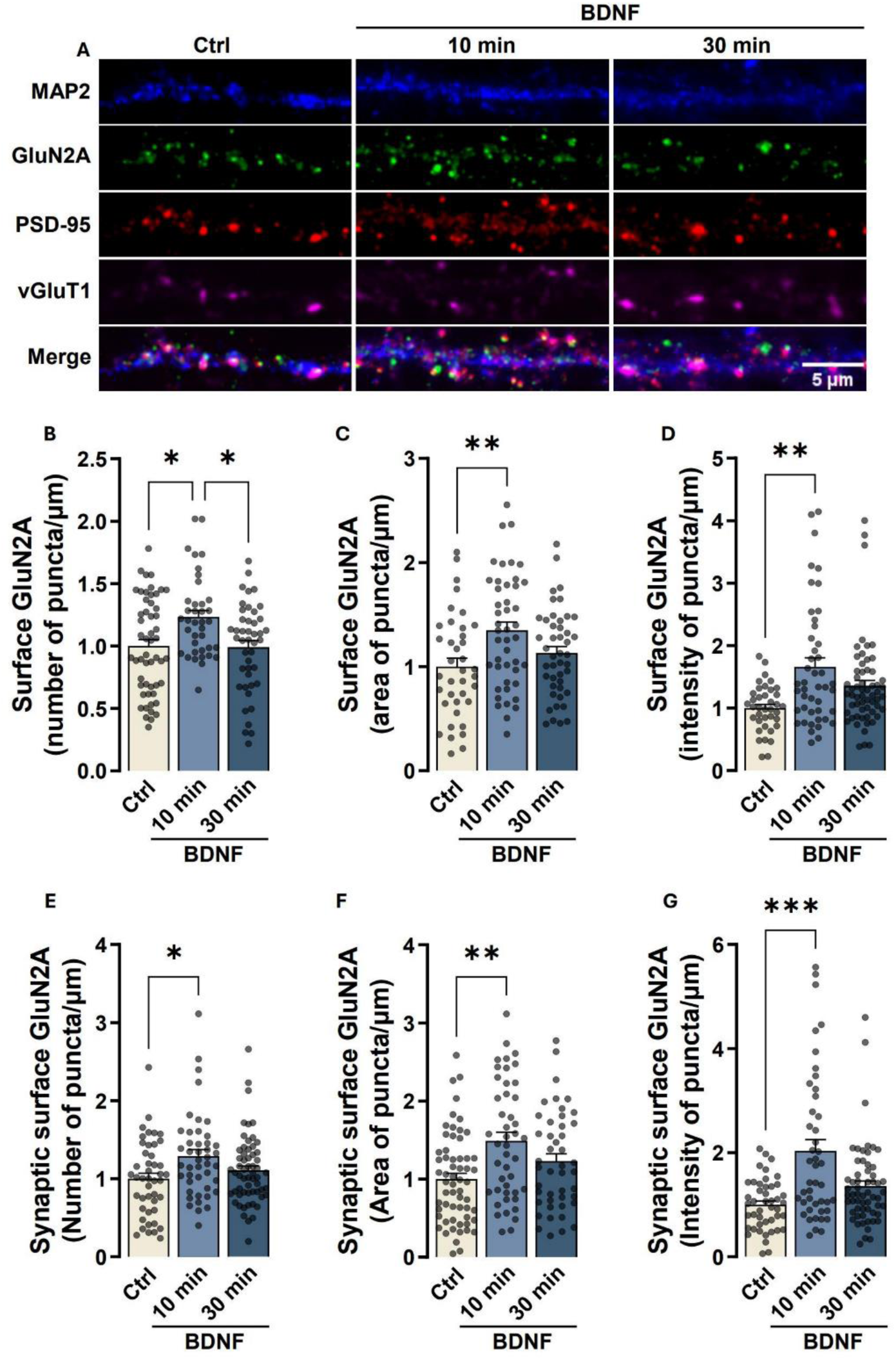
Stimulation of hippocampal neurons with BDNF for 10 min increases the synaptic expression of GluN2A-containing NMDAR. (**A**) Representative images of cultured hippocampal neurons (DIV 14 - 15) stimulated with BDNF (50 ng/ml for 10 or 30 min), as indicated. Neurons were then live-immunoassayed for GluN2A using an antibody against an extracellular epitope in the GluN2A N-terminus, fixed, and further immunoassayed for PSD-95, vGluT1, and MAP2. Scale bar, 5 μm. Images illustrated in (**A**) were analyzed for the total number (**B**), area (**C**), and intensity (**D**) of surface GluN2A puncta per dendritic length. Synaptic (PSD-95- and vGluT1-colocalized) surface GluN2A number (**E**), area (**F**), and intensity (**G**) of puncta per density of excitatory synapses (number of puncta PSD-95–vGluT1 colocalized per µm), were also analyzed. Data are normalized to the mean of the control and are the means ± SEM of 43-45 cells per condition, from at least three independent experiments performed in different preparations. *p < 0.05, **p < 0.01, ***p <0.001 by Kruskal-Wallis’s test and Dunn’s multiple comparisons test.

In complementary experiments, we assessed the time-dependent effects of BDNF on the synaptic surface accumulation of GluN2A in rat hippocampal synaptoneurosomes (Fig. 2), which are subcellular fractions containing the pre- and postsynaptic components. Synaptoneurosomes were stimulated with 50 ng/mL BDNF for 10, 30, and 60 min before live immunostaining with an antibody against the extracellular N-terminus of GluN2A, which allows labeling surface receptors (Fig. 2A). Synaptic surface GluN2A-NMDAR were identified by colocalization with the pre-and post-synaptic markers vesicular glutamate transporter 1 (vGluT1) and PSD-95, respectively. The results were analyzed for the integrated density (Fig. 2B) and area (Fig. 2C) of GluN2A clusters immunoreactivity that colocalized with the synaptic markers. Incubation with BDNF for 10, 30, and 60 min upregulated the integrated density (Fig. 2B) and cluster area (Fig. 2C) of GluN2A immunoreactivity in hippocampal synaptoneurosomes.

**Figure 2.**
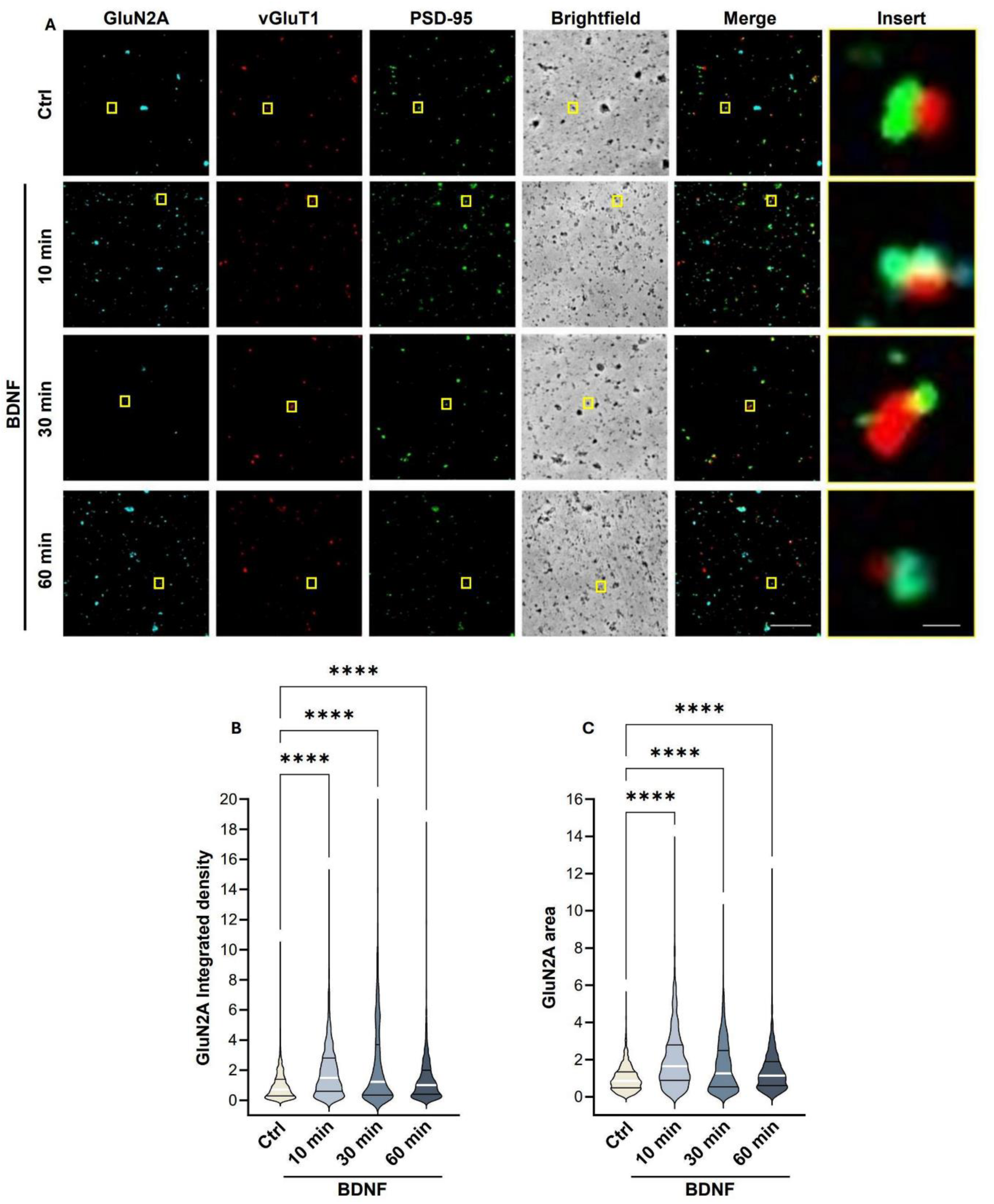
BDNF-induced increase in the expression of GluN2A-containing NMDAR in hippocampal synaptoneurosomes. (**A**) Representative images of hippocampal synaptoneurosomes (prepared from adult Sprague-Dawley rats 6-8 weeks old) incubated with BDNF (50 ng/mL). Synaptoneurosomes were immunoassayed for GluN2A using an antibody against an extracellular epitope in the GluN2A N-terminus and immunoassayed for vGluT1, PSD-95. Merge scale bar, 10 μm. Insert scale bar, 0.5 um. Images illustrated in (**A**) were analyzed for the GluN2A integrated density (**B**) and area (**C**). Data are the means ± SEM of 779 - 840 synaptoneurosomes per condition, in at least three independent experiments performed in different preparations. ****p < 0.0001, by Kruskal-Wallis’s test and Dunn’s multiple comparisons test.

Taken together, these results indicate that BDNF rapidly enhances the synaptic surface expression of GluN2A-NMDAR both in cultured hippocampal neurons and in hippocampal synapses from adult rats, although this effect is transient in cultured neurons and more sustained in synaptoneurosomes derived from mature hippocampal tissue.

### BDNF increases synaptic expression of GluN2A-NMDAR in a protein synthesis–dependent manner

Previous studies showed that the effects of BDNF on the synaptic delivery of NMDAR are dependent on protein synthesis (Leal, Comprido et al. 2017, Afonso, De Luca et al. 2019). However, it remains to be determined whether translation activity is specifically required for BDNF-induced synaptic surface accumulation of GluN2A-NMDAR. To address this question, we tested the effect of the translation inhibitor cycloheximide (CHX; 50 µg/ml) on BDNF-induced alterations in the surface distribution of GluN2A in cultured hippocampal neurons. In these experiments, the cells were stimulated with BDNF for 10 min, which leads to a significant increase in the total and synaptic surface distribution of GluN2A-NMDAR (Figures 1 and 3). Pre-incubation with CHX completely abrogated the effects of BDNF on the total number (Fig. 3B), area (Fig. 3C), and intensity (Fig. 3D) of surface GluN2A dendritic puncta, as well as on the synaptic expression of the receptors (Fig. 3E, F, G). Together, this evidence shows that the BDNF-induced upregulation in total and synaptic expression of GluN2A-NMDAR is a protein synthesis-dependent mechanism.

**Figure 3.**
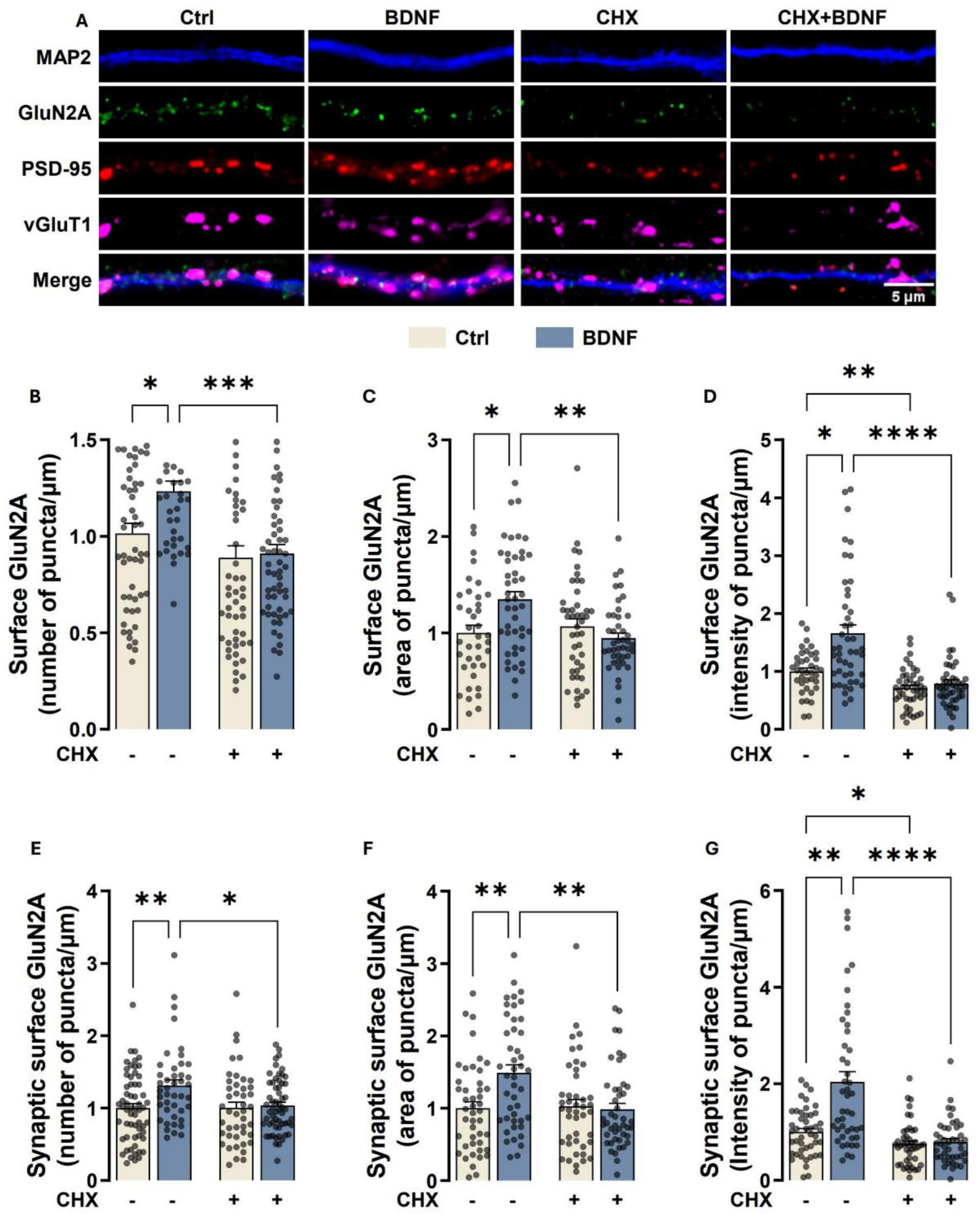
BDNF-induced increase in the synaptic expression of GluN2A-containing NMDAR is protein syntheses dependent. (**A**) Representative images of hippocampal neurons (DIV 14 - 15) pre-incubated with CHX (50 µg/ml) or vehicle (DMSO; 1:1000 dilution for 40 min) and then either maintained under the same conditions or stimulated with BDNF (50 ng/ml for 10 min), as indicated. Neurons were live-immunoassayed for GluN2A using an antibody against an extracellular epitope in the GluN2A N-terminus, fixed, and then further immunoassayed for PSD-95, vGluT1, and MAP2. Scale bar, 5 μm. Images illustrated in (**A**) were analyzed for the total number (**B**), area (**C**), and intensity (**D**) of surface GluN2A puncta per dendritic length. Synaptic (PSD-95- and vGluT1-colocalized) surface GluN2A number (**E**), area (**F**), and intensity (**G**) of puncta per µm of excitatory synapses (number of puncta PSD-95–vGluT1 colocalized per µm), were also analyzed. Data are normalized to the mean of DMSO control and are the means ± SEM of 58 - 60 cells per condition, in at least three independent experiments performed in different preparations. *p < 0.05, **p < 0.01, ***p < 0.01, ****p < 0.0001 by Kruskal-Wallis’s test and Dunn’s multiple comparisons test.

### BDNF-mediated GluN2A upregulation in hippocampal neurons requires hnRNPK-Pyk2-PKC signaling

Previous evidence from our laboratory showed that the RNA-binding protein hnRNPK mediates the effect of BDNF on dendritic mRNA metabolism and regulates synaptic NMDA receptors in cultured hippocampal neurons (Leal, Comprido et al. 2017). Moreover, Pyk2, a non-receptor tyrosine kinase highly enriched in forebrain neurons, plays an important role in regulating GluN2A-NMDAR (Montalban, Al-Massadi et al. 2019, de Pins, Mendes et al. 2021). CAKβ/Pyk2 has been reported to be activated by various stimuli, possibly through stimulation of protein kinase C (PKC) (Lev, Moreno et al. 1995). In addition, protein Kinase C (PKC), Pyk2, and NMDA receptors are interconnected components of a signaling cascade that plays a crucial role in synaptic plasticity and neuronal function (Huang, Lu et al. 2001, Bartos, Ulrich et al. 2010). Based on this evidence, we hypothesized that these signaling players may play a key role on the regulation of the GluN2A-containing NMDAR on the synaptic surface mediated by BDNF.

The role of hnRNPK in BDNF-induced increase in the surface abundance of GluN2A-NMDAR was investigated in cultured hippocampal neurons using short hairpin RNAs (shRNAs) to knock down hnRNPK (sh-hnRNPK). This shRNA was previously shown to downregulate hnRNPK protein levels in cultured hippocampal neurons (Leal, Comprido et al. 2017). Knockdown of hnRNPK abrogated the effects of BDNF on the total surface expression of GluN2A subunit in the dendritic compartment, as determined by live immunostaining with an antibody against an extracellular epitope in the N-terminal region of the protein and by colocalization with the dendritic marker MAP2 (Fig. 4B, C, D). We then investigated whether hnRNPK also plays a role in the BDNF-induced up-regulation of GluN2A surface expression at the synaptic compartment, as determined by colocalization with the pre- and post-synaptic marker vGluT1 and PSD-95, respectively. BDNF significantly increased the number (Fig. 4E), area (Fig. 4F), and intensity (Fig. 4G) of synaptic surface GluN2A puncta in neurons transfected with the control shRNA, and knockdown of hnRNPK abolished these effects of the neurotrophin (Fig. 4E-G).

**Figure 4.**
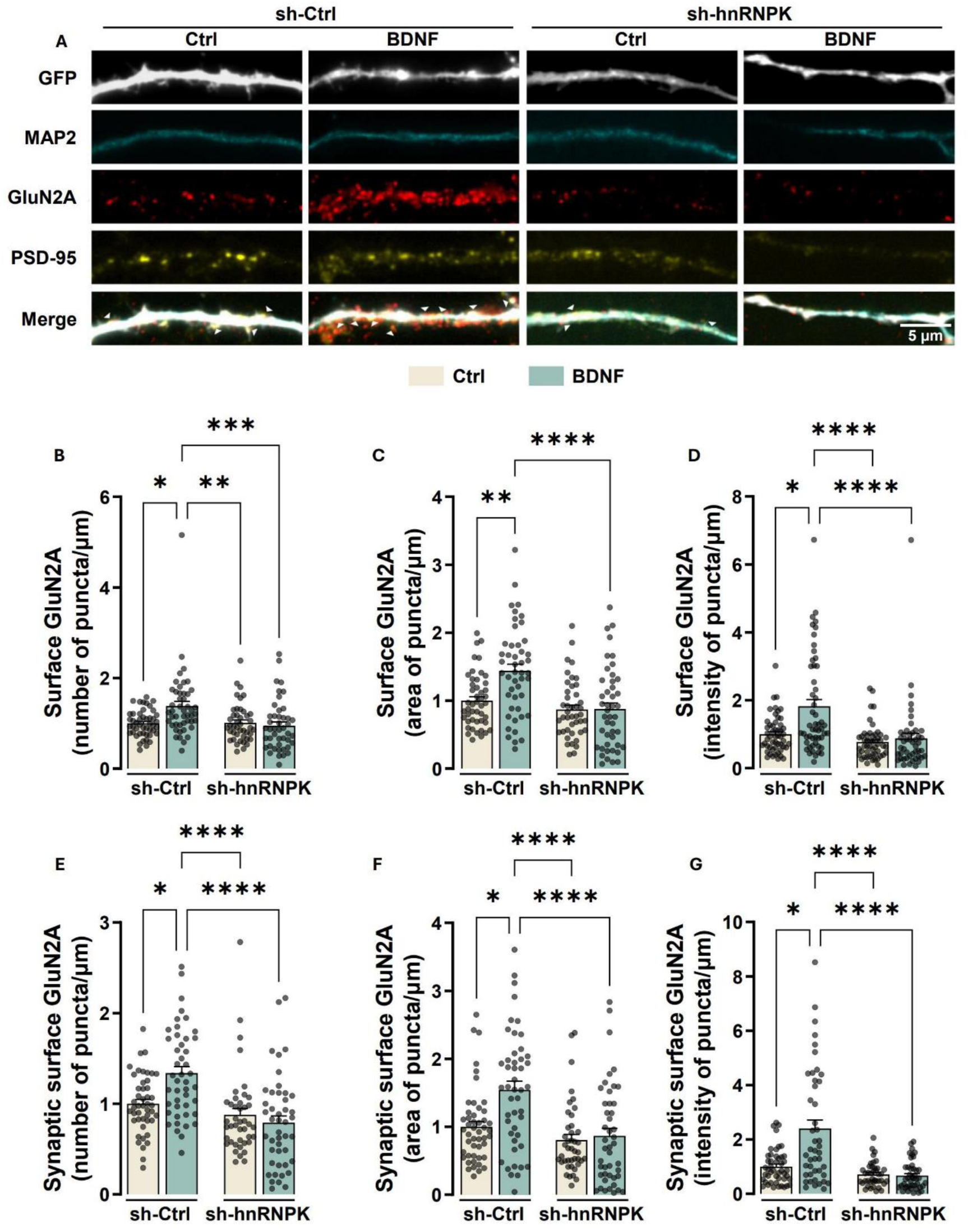
BDNF-induced increase in the synaptic surface expression of NMDAR-containing GluN2A subunits requires hnRNPK. (A) Representative images of rat hippocampal neurons transfected with short hairpin control (sh-Ctrl) or with a hnRNPK-targeted short hairpin (sh-hnRNPK) RNA at DIV 12. At DIV 15, cultures were then either maintained under the same conditions or stimulated with BDNF (50 ng/ml for 10 min) as indicated. Neurons were live-immunoassayed for GluN2A using an antibody against an extracellular epitope in the GluN2A N-terminus, fixed, and then further immunoassayed for PSD-95 and MAP2. Scale bar, 5 μm. Images illustrated in (**A**) were analyzed for the total number (**B**), area (**C**), and intensity (**D**) of surface GluN2A puncta per dendritic length. Synaptic (PSD-95-colocalized) surface GluN2A number (**E**), area (**F**), and intensity (**G**) of puncta per µm of excitatory synapses (number of puncta PSD-95-vGluT1 colocalized per µm), were also analyzed. Data are normalized to the mean of sh-Ctrl and are the means ± SEM of 44 - 47 cells per condition, in at least three independent experiments performed in different preparations. *p < 0.05, **p < 0.01, ***p < 0.001, ****p < 0.0001 by Kruskal-Wallis’s test and Dunn’s multiple comparisons test.

The Pyk2 mRNA was among the transcripts that co-immunoprecipitated with hnRNPK in studies performed in cultured hippocampal neurons, and this interaction decreased following stimulation with BDNF (Leal et al., 2017). Therefore, we investigated whether Pyk2 also plays a role in the BDNF-induced up-regulation of GluN2A surface expression at the synaptic compartment, as determined by colocalization with the postsynaptic marker PSD95. The role of Pyk2 in BDNF-induced increase in the surface abundance of GluN2A-NMDAR was investigated using short hairpin RNAs (shRNAs) to knock down Pyk2 (sh-Pyk2). This shRNA was previously shown to abrogate the effects mediated by Pyk2 in cultured hippocampal neurons (Afonso, De Luca et al. 2019). Knockdown of Pyk2 abrogated the effects of BDNF on the total surface expression of GluN2A subunit in the dendritic compartment, as determined by live immunostaining with an antibody against an extracellular epitope in the N-terminal region of the protein and by colocalization with the dendritic marker MAP2 (Fig. 5A-D). We then investigated whether Pyk2 also plays a role in the BDNF-induced up-regulation of GluN2A surface expression at the synaptic compartment, as determined by colocalization with the post-synaptic marker PSD-95. BDNF significantly increased the number (Fig. 5E), area (Fig. 5F) and intensity (Fig. 5G) of synaptic surface GluN2A puncta in neurons transfected with the control shRNA, whereas knockdown of Pyk2 abolished these effects of the neurotrophin. Interestingly, the levels of total and synaptic surface GluN2A intensity were significantly decreased when Pyk2 was downregulated (see Figure 5D and G). However, this reduction in GluN2A intensity was completely antagonized by stimulation with BDNF (Figure 5D and G).

**Figure 5.**
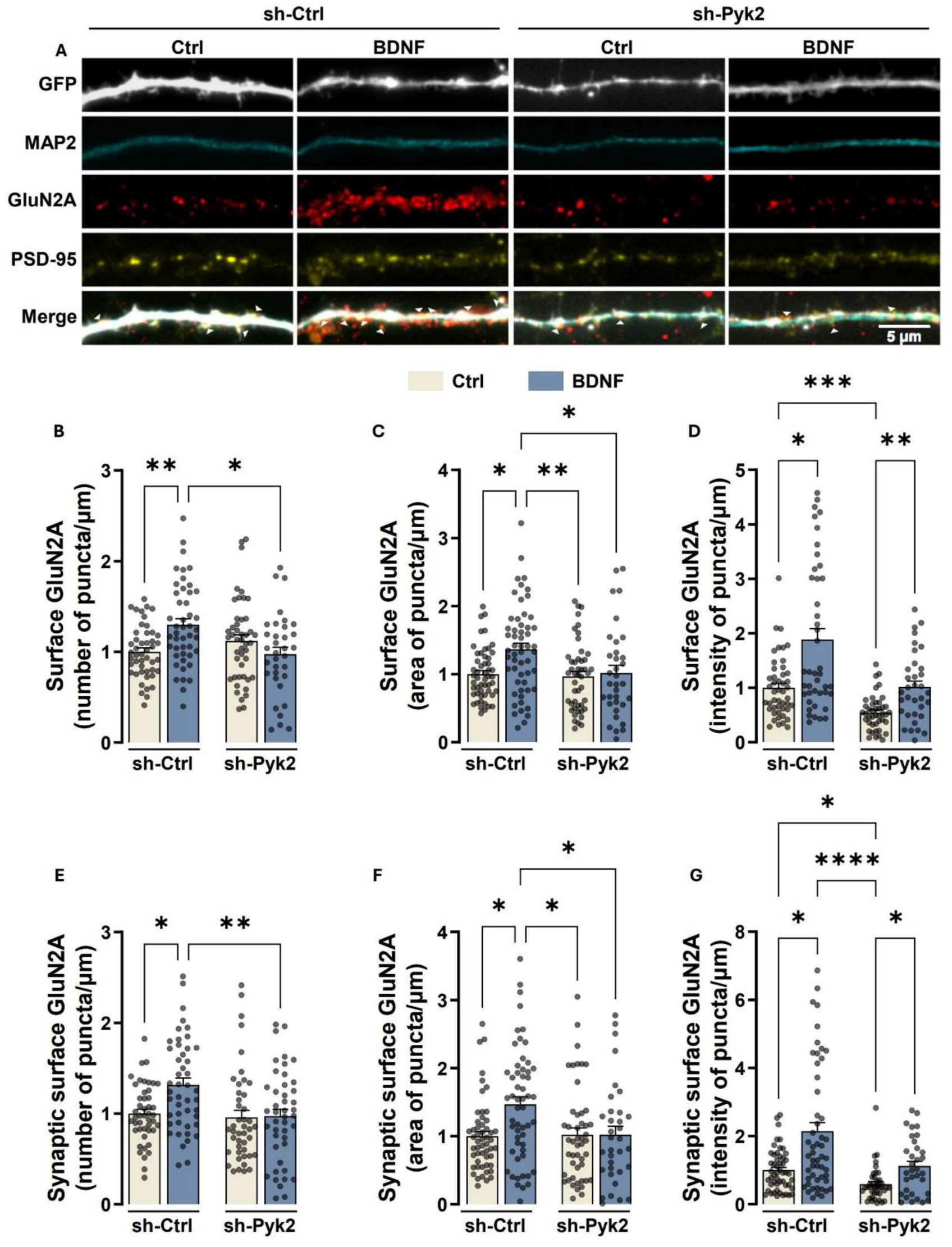
BDNF-induced increase in the synaptic surface expression of NMDAR-containing GluN2A subunits requires Pyk2. (**A**) Representative images of rat hippocampal neurons transfected with short hairpin control (sh-Ctrl) or with a Pyk2-targeted short hairpin (sh-Pyk2) RNA at DIV 12. At DIV 15, cultures were then either maintained under the same conditions or stimulated with BDNF (50 ng/ml for 10 min) as indicated. Neurons were live-immunoassayed for GluN2A using an antibody against an extracellular epitope in the GluN2A N-terminus, fixed, and then further immunoassayed for PSD-95 and MAP2. Scale bar, 5 μm. Images illustrated in (**A**) were analyzed for the total number (**B**), area (**C**), and intensity (**D**) of surface GluN2A puncta per dendritic length. Synaptic (PSD-95-colocalized) surface GluN2A number (**E**), area (**F**), and intensity (**G**) of puncta per µm of excitatory synapses (number of puncta PSD-95-vGluT1 colocalized per µm), were also analyzed. Data are normalized to the mean of sh-Ctrl and are the means ± SEM of 44 - 47 cells per condition, in at least three independent experiments performed in different preparations. *p < 0.05, **p < 0.01, ***p < 0.001, ****p < 0.0001 by Kruskal-Wallis’s test and Dunn’s multiple comparisons test.

To determine whether the BDNF-induced increase in GluN2A surface expression is mediated by the activation and phosphorylation of Pyk2, we examined the effect of BDNF in hippocampal neurons transfected with either a phospho-mutant form of Pyk2 (Pyk2-Y402F), which lacks kinase activity, or the wild-type form of Pyk2 (Pyk2-WT). As a control, hippocampal neurons were transfected with a vector expressing Flag (Flag). Live immunostaining using an antibody targeting an extracellular epitope at the N-terminus of GluN2A revealed that the BDNF-induced upregulation of GluN2A surface expression was abolished in neurons expressing Pyk2-Y402F (Fig. 6A-D), as well as on the synaptic expression of the receptors (Fig. 6A, E-G). Notably, overexpression of Pyk2-Y402F led to a significant reduction in the area of puncta of basal GluN2A surface expression under resting conditions (Fig. 6C). suggesting that Pyk2 phosphorylation might be required for maintaining the spatial distribution of surface GluN2A levels under resting conditions. Conversely, transfection with Pyk2-WT mimicked the effects of BDNF, leading to an increase in the dendritic surface expression of GluN2A-NMDAR (Fig. 6A-D), as well as on the synaptic expression of the receptors (Fig. 6E, F, G), reinforcing the role of Pyk2 activity in regulating GluN2A trafficking. These findings collectively indicate that the phosphorylation of Pyk2 at Y402 is essential for both the basal maintenance and BDNF-induced upregulation of GluN2A surface expression in cultured hippocampal neurons.

**Figure 6.**
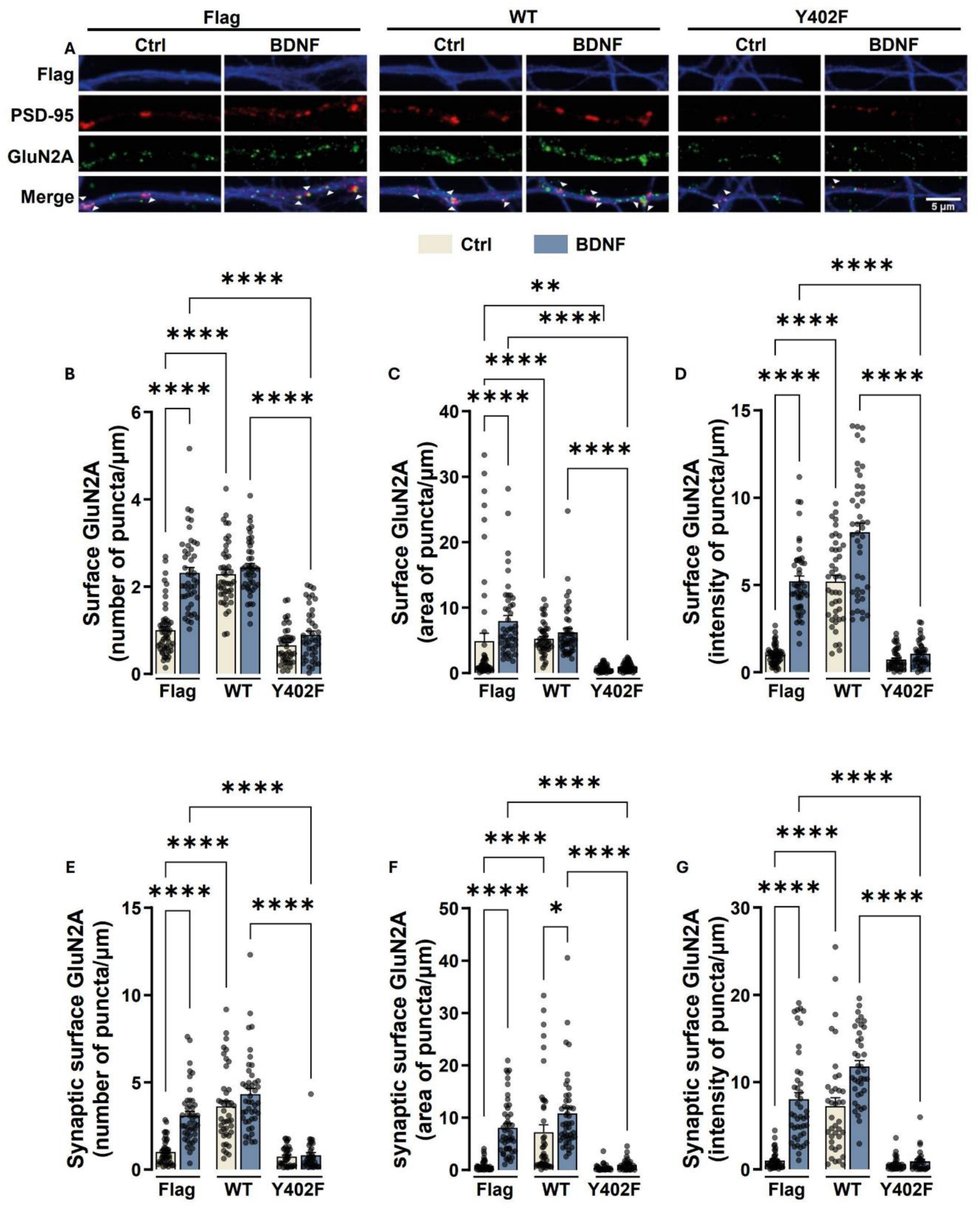
BDNF-induced increase in synaptic expression of GluN2A-containing NMDAR is mediated by activation of Pyk2. (A) Representative images of hippocampal neurons that were transfected with wild-type (WT) or phospho-mutant, kinase-deficient Pyk2 (Y402F) at DIV 12, and at DIV 15 were maintained under control conditions or stimulated with BDNF (50 ng/ml) for 10 min, live-immunostained for GluN2A (using an antibody against an extracellular epitope in the GluN2A N terminus), fixed and permeabilized, and further immunostained for Flag (transfection marker) and MAP2. Scale bar, 5 µm. Images illustrated in (**A**) were analyzed for the total number (**B**), area (**C**), and intensity (**D**) of surface GluN2A puncta per dendritic length. Synaptic (PSD-95-colocalized) surface GluN2A number (**E**), area (**F**), and intensity (**G**) of puncta per µm of excitatory synapses (number of puncta PSD-95-vGluT1 colocalized per µm), were also analyzed. Data are normalized to the mean of the empty vector control (Flag) and are means ± SEM of 45 - 34 cells per condition, in at least three independent experiments performed in different preparations. *p < 0.05; **p < 0.01, ****p < 0.0001 by Kruskal-Wallis’s test and Dunn’s multiple comparisons test.

Previous studies have demonstrated that protein kinase C (PKC) activation can trigger Pyk2 activity (de Pins, Mendes et al. 2021), and PKC itself is activated following TrkB-mediated stimulation of phospholipase C-γ (Barry and McGinty 2017). This suggests a potential pathway wherein BDNF-TrkB signaling leads to PKC stimulation, which in turn activates Pyk2, culminating in the upregulation of GluN2A synaptic surface expression. Given the established role of PKC in mediating BDNF-induced regulation of Pyk2, we investigated whether this signaling pathway similarly mediates the neurotrophin-driven synaptic surface expression of GluN2A-NMDAR. To this end, we assessed the synaptic surface expression of GluN2A-NMDAR in cultured hippocampal neurons subjected to BDNF stimulation for 10 min or maintained under control conditions. Quantitative image analysis was performed to evaluate the number, intensity, and spatial distribution of total and synaptic GluN2A puncta per dendritic length.

Pharmacological inhibition of PKC with GÖ 6983 abolished the BDNF-mediated enhancement of the total surface GluN2A puncta number (Fig. 7A, B), as well as their area (Fig. 7A, C) and intensity (Fig. 7A, D). Similarly, PKC inhibition prevented the BDNF-induced increase in the number of synaptic surface GluN2A puncta (Fig. 7A, E), along with their respective area (Fig. 7A, F) and intensity (Fig. 7A, G). These findings demonstrate that PKC is a pivotal mediator of BDNF-induced modulation of synaptic GluN2A-NMDAR expression, underscoring its crucial role in neurotrophin-dependent synaptic plasticity. In summary, our findings uncover a crucial signaling axis wherein BDNF-mediated TrkB activation engages PKC and Pyk2 to drive the synaptic enrichment of GluN2A-containing NMDARs. Furthermore, the involvement of hnRNPK suggests a potential regulatory checkpoint that fine-tunes this cascade, linking neurotrophic signaling to synaptic plasticity.

**Figure 7.**
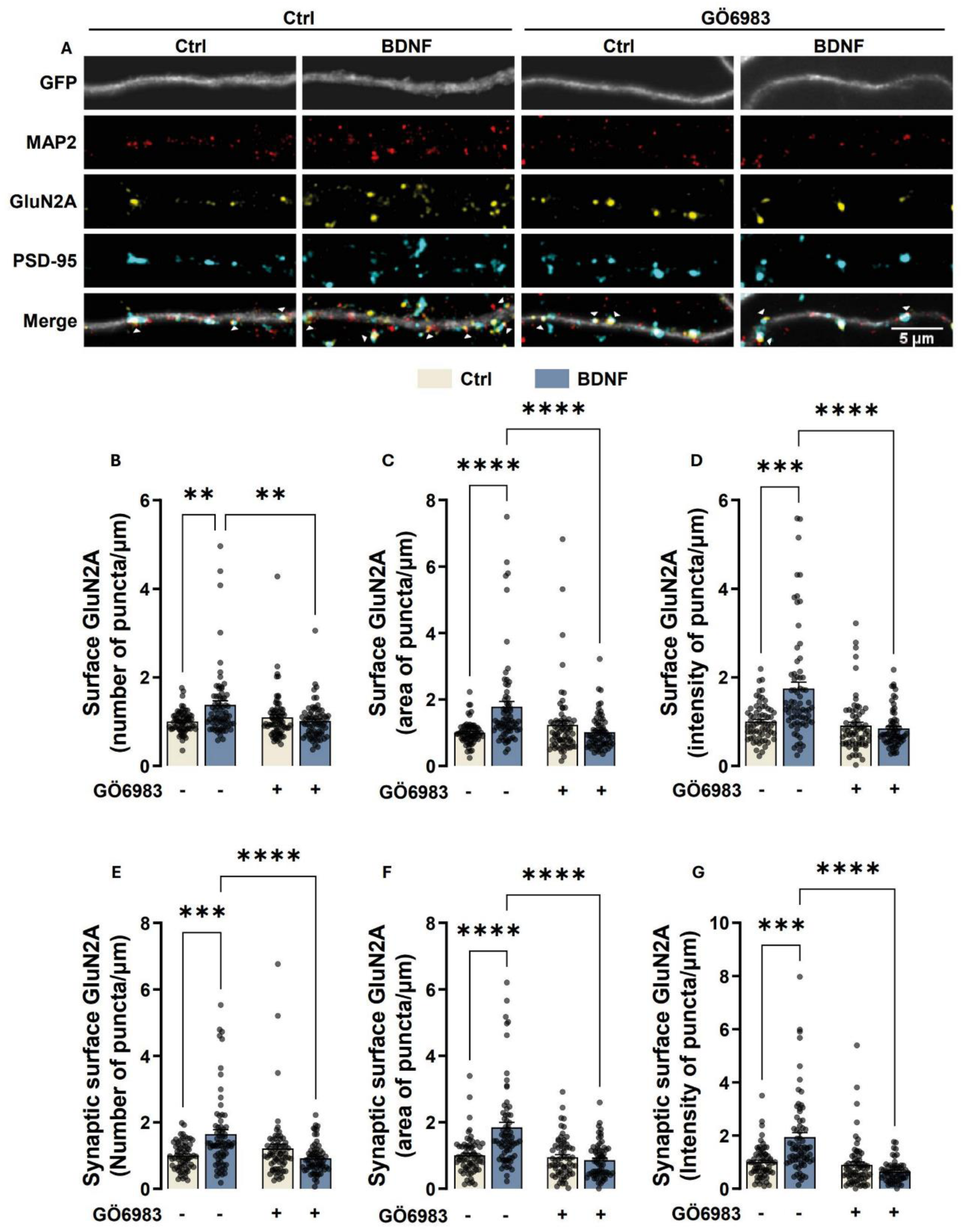
BDNF-induced increase in the synaptic expression of GluN2A-containing NMDAR is sensitive to PKC inhibition. (A) Representative images of hippocampal neurons (DIV 14 - 15) pre-incubated with GÖ 6983 (100 nM) or vehicle (DMSO; 1:1000 dilution for 40 min) and then either maintained under the same conditions or stimulated with BDNF (50 ng/ml for 10 min), as indicated. Neurons were live-immunoassayed for GluN2A using an antibody against an extracellular epitope in the GluN2A N-terminus, fixed, and then further immunoassayed for PSD-95, vGluT1, and MAP2. Scale bar, 5 μm. Images illustrated in (**A**) were analyzed for the total number (**B**), area (**C**), and intensity (**D**) of surface GluN2A puncta per dendritic length. Synaptic (PSD-95-colocalized) surface GluN2A number (**E**), area (**F**), and intensity (**G**) of puncta per µm of excitatory synapses (number of puncta PSD-95-vGluT1 colocalized per µm), were also analyzed. Data are normalized to the mean of the DMSO control and are the means ± SEM of 67 - 70 cells per condition, in at least three independent experiments performed in different preparations. **p < 0.01, ***p < 0.001 ** **p < 0.0001 by Kruskal-Wallis’s test and Dunn’s multiple comparisons test.

### TrkB signaling regulates synaptic GluN2A expression in status epilepticus

BDNF-TrkB signaling has been shown to play a key role in epileptogenesis (Gu, Huang et al. 2015, Huang, He et al. 2019, Krishnamurthy, Huang et al. 2019). In addition, an upregulation in TrkB receptor activity was observed in hippocampal synaptoneurosomes isolated from rats subjected to pilocarpine-induced status epilepticus (SE) (De Luca, Mele et al. 2025). To determine whether BDNF-TrkB signaling is coupled to the regulation of the synaptic expression of GluN2A in SE, we analyzed the synaptic surface abundance of this NMDAR subunit in synaptoneurosomes from rats subjected to the pilocarpine model of temporal lobe epilepsy (TLE) under control conditions and after treatment with ANA-12, a TrkB receptor inhibitor (Fig. 8A). The surface expression of GluN2A was assessed as described in the experiments illustrated in Fig. 1 and representative images are shown in Fig. 8B. The integrated density and area of GluN2B staining was increased in animals treated with pilocarpine, and this effect was prevented by administration of ANA-12 (Fig. 8B-D). In addition, administration of ANA-12 under control conditions did not change GluN2A staining in hippocampal synaptoneurosomes. These results indicate that inhibition of TrkB receptor effectively blocks the downstream signaling pathways that are activated under conditions of SE.

**Figure 8.**
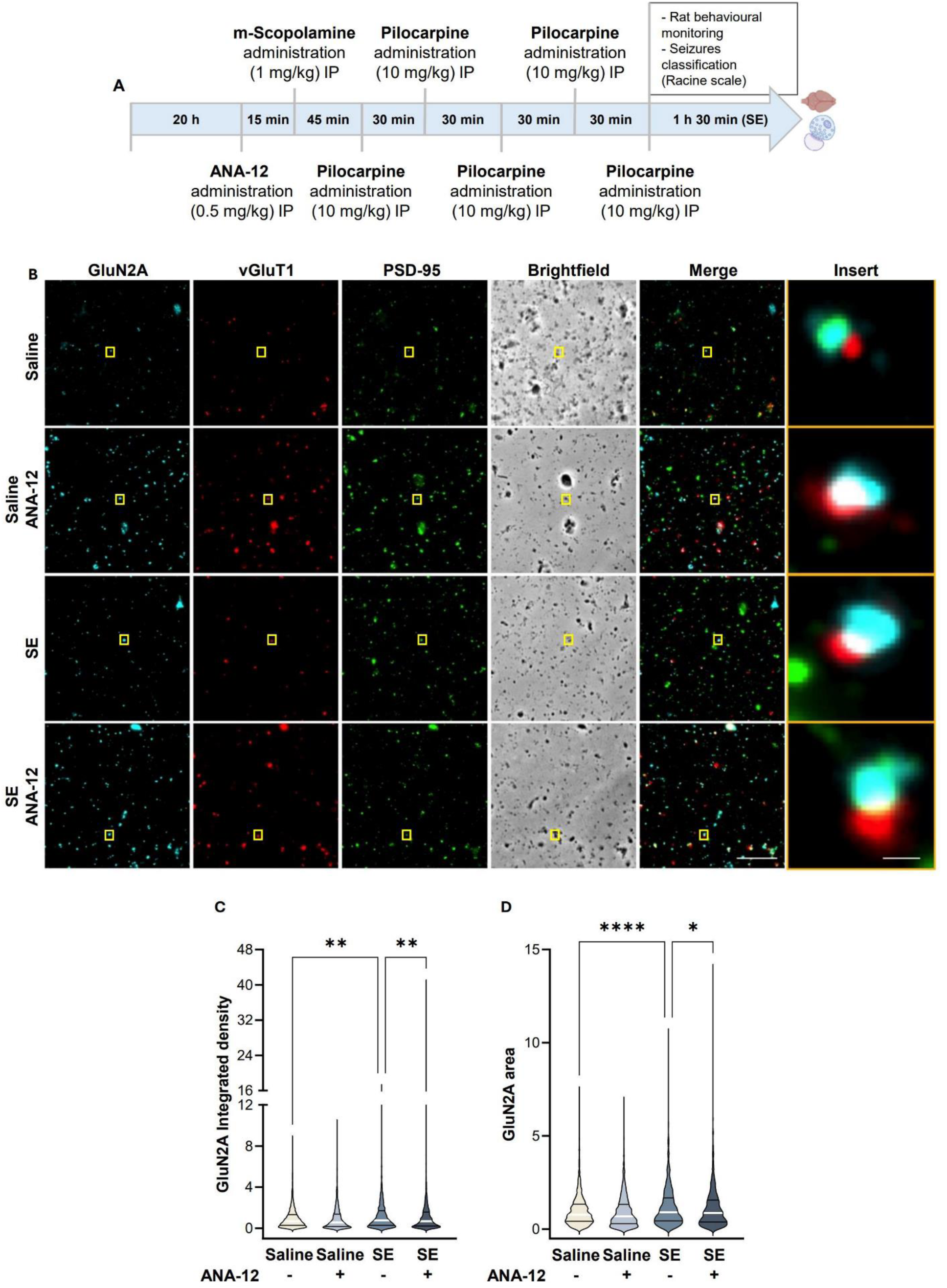
The increase in the expression of GluN2A-containing NMDAR in hippocampal synaptoneurosomes during status epilepticus requires TrkB signaling. (**A**) Experimental design for the lithium-pilocarpine model of Status Epilepticus. (**B**) Representative images of hippocampal synaptoneurosomes (prepared from adult rats treated with saline, saline and ANA-12, Pilocarpine or Pilocarpine and ANA-12). Synaptoneurosomes were live-immunoassayed for GluN2A using an antibody against an extracellular epitope in the GluN2A N-terminus and immunoassayed for vGluT1 and PSD-95. Merge scale bar, 10 μm. Insert scale bar, 0.5 µm. Images illustrated in (**B**) were analyzed for the GluN2A integrated density (**C**) and GluN2A area (**D**). Data are the means ± SEM of 1592 - 2008 synaptoneurosomes per condition, from at least four animals for each experimental condition. *p < 0.05, **p < 0.01, ****p < 0.0001, as determined by Kruskal Wallis’s test and Dunn’s multiple comparisons test.

### Endogenous BDNF and GluN2A-NMDAR contributed to the alterations in neuronal network activity induced by neuronal hyperexcitability

To further investigate the functional impact of BDNF-TrkB signalling on neuronal network dynamics and the role of GluN2A in this context, we conducted microelectrode array (MEA) recordings to assess evoked activity in cultured hippocampal neurons. Recordings were obtained under control conditions (Baseline, BL) and after treatment with a cocktail comprising bicuculline, 4-aminopyridine (4-AP), and glycine, to induce epileptiform synchronization of the hippocampal neurons (bicuculline, 4-AP, and glycine) (Avoli, de Curtis et al. 2016). The first baseline recording (BL1) was collected before pharmacological interventions, and the second recording (BL2) was performed after 30 min of preincubation with TrkB-Fc (1 µg/mL) or TCN 213 (10 µM), or the respective vehicles. Stimulation with bicuculline and 4-AP together with glycine significantly increased burst frequency (Fig. 9A, B, and C) and network burst frequency (Fig. 9A, D and E), reflecting enhanced network excitability and synchronization. To assess the role of TrkB signalling under these conditions, we used TrkB-Fc, a BDNF scavenger, which led to a reduction in burst frequency (Fig. 9A, B) and network burst frequency (Fig. 9A, D) compared to control conditions. These results indicate that BDNF/TrkB signalling plays a critical role in maintaining heightened network activity during synaptic excitatory stimulus.

**Figure 9.**
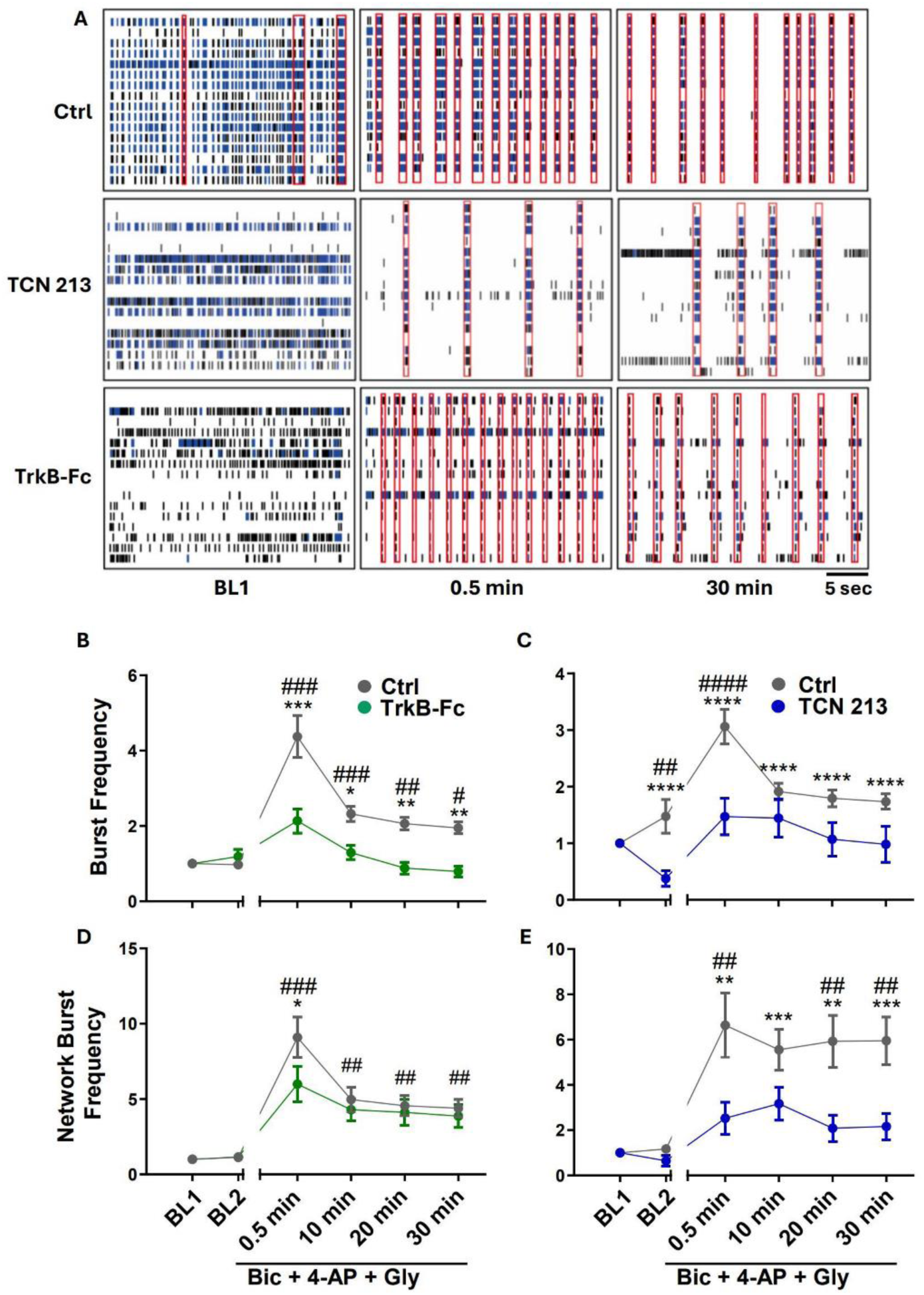
Endogenous BDNF and GluN2A-containing NMDA receptors contributed to depolarization-induced increase in neuronal network activity in cultured hippocampal neurons. (**A**) Representative raster plots of cultured hippocampal neurons showing 30 sec of electrical activity. Basal spontaneous activity of neuronal cells cultured on MEA plates was recorded at 16 DIV (baseline recording, 3 minutes). (**B** - **J**) After baseline recording (BL1 and BL2), a chemical synaptic stimulation was performed (2.5 mM 4-AP, 50 μM Bicuculline, 10 μM Glycine), and neuronal activity was recorded after stimulation every 10 min, for 30 min. In the experiments performed in the presence of TCN 213 (10 μM) or TrkB-Fc (1 μg/mL), the inhibitors were added to the medium 30 minutes before synaptic stimulation, and a recording was performed at the end of the 30 min preincubation (BL 2). The parameters extracted from the analysis of this study were: (I) the burst frequency (BF) (**B, C**), and the network burst frequency (NBF) (**D, E**). Statistical analysis for the BF and NBF was performed by Two-way analysis of variance (ANOVA) followed by the Bonferroni comparisons test. (^#^p < 0.05, ^##^p < 0.01, ^####^p < 0.0001 compared to the baseline, *p < 0.05, **p < 0.01, ****p < 0.0001, compared to the control condition).

Furthermore, to determine the role of GluN2A in the depolarization-induced responses in network activity, we incubated cells with TCN 213, a selective GluN2A inhibitor. This treatment resulted in a significant decrease in both burst and network burst frequency, suggesting a pivotal role of GluN2A-NMDAR in the neuronal network activity alterations induced by excitatory stimulation. The inhibition of the responses induced by bicuculline + 4-AP + glycine upon blockage of the BDNF signalling and GluN2A-NMDAR suggests that the latter population of receptors mediates the effects of BDNF in the neuronal network activity.

## Discussion

Different processes involving plastic changes at hippocampal glutamatergic synapses, including LTP and epileptogenesis, have been associated with an upregulation in BDNF signaling (Korte, Carroll et al. 1995, Carvalho, Caldeira et al. 2008, Leal, Comprido et al. 2014, Gu, Huang et al. 2015, Huang, He et al. 2019, Krishnamurthy, Huang et al. 2019, Mohandas, Rishikesh et al. 2025) and increased activity of GluN2A-NMDAR (Chen, He et al. 2007, Franchini, Carrano et al. 2020). In this work, we show that BDNF mediates the upregulation in the synaptic expression of GluN2A in the hippocampus under conditions characterized by an activity-dependent increase of glutamatergic activity. This model is supported by several different pieces of evidence: (i) BDNF stimulation evoked a time-dependent increase in the surface expression of GluN2A in hippocampal synaptoneurosomes; (ii) incubation of cultured hippocampal neurons with BDNF upregulated the synaptic expression of GluN2A by a mechanism dependent of protein synthesis and involving the contribution of the RNA binding protein hnRNP K; (iii) inhibition of TrkB receptors with ANA-12 decreased the surface accumulation of GluN2A in hippocampal synaptoneurosomes isolated from rats subjected to the pilocarpine model of temporal lobe epilepsy; (iv) BDNF signaling and GluN2A-NMDAR mediated the depolarization-induced increase in neuronal synchronization in hippocampal cultures.

In this study, we found that a 10-min incubation with BDNF increases the synaptic surface expression of GluN2A-NMDAR in both cultured hippocampal neurons and hippocampal synaptoneurosomes, as indicated by their colocalization with the synaptic markers vGluT1 and PSD-95. Previous biotinylation studies have shown that BDNF enhances the total surface expression of GluN2A only after 24 h of incubation (Caldeira, Melo et al. 2007), suggesting that the rapid synaptic accumulation observed here results from a redistribution of receptors already present at the cell surface. Alternatively, it may depend on a pool of receptors that are delivered to the cell surface or undergo reduced internalization in the perisynaptic region and are therefore not detected in the biotinylation assay. The incubation period required to induce the synaptic accumulation of GluN2A was shorter than the time previously reported for BDNF to induce the synaptic accumulation of GluN2B-NMDAR in both preparations (De Luca, Mele et al. 2025). Protein-protein interactions between the C-terminus of GluN subunits and MAGUKs favor the anchoring and stabilization of diffusing NMDAR at the postsynaptic density (Tahiri, Corti et al. 2025), and GluN2B-NMDAR are less trapped at the synapse than GluN2A-NMDAR (Groc, Heine et al. 2006). This difference may account, at least in part, for the slower rate of synaptic accumulation of GluN2A compared with GluN2B following BDNF stimulation. The faster rate of synaptic accumulation of GluN2A-NMDA following BDNF stimulation will change the ratio GluN2A/GluN2N-NMDA, which directly influences activity-dependent long-term changes in synaptic strength. An increase in the contribution of GluN2A-NMDAR is typically associated with an increase in the threshold for further potentiation (Barria and Malinow 2005, Gardoni, Mauceri et al. 2009, Xu, Chen et al. 2009).

The BDNF-induced synaptic accumulation of GluN2A-NMDAR was blocked by the protein synthesis inhibitor cycloheximide and by downregulation of the RNA-binding protein hnRNP K. This latter finding is consistent with previous reports indicating that hnRNP K plays a key role in mediating the effects of BDNF on dendritic mRNA metabolism (Leal, Comprido et al. 2017). Studies in cultured hippocampal neurons have shown that BDNF-TrkB signaling promotes the dendritic accumulation of hnRNP K and triggers the release of numerous transcripts encoding synaptic proteins (Leal, Comprido et al. 2017), supporting a key role for this RNA-binding protein in BDNF-mediated forms of synaptic plasticity (Korte, Carroll et al. 1995, Kang and Schuman 1996, Panja, Kenney et al. 2014, Santos, Mele et al. 2015, Costa, Martins et al. 2022, De Luca, Mele et al. 2025). Several lines of evidence suggest that hnRNP K-dependent, BDNF-induced local synthesis of Pyk2 may be a key mechanism underlying the neurotrophin-driven upregulation of synaptic GluN2A: (i) the mRNA for Pyk2 co-immunoprecipitated with hnRNP K, and a decrease in this interaction was observed following stimulation of cultured hippocampal neurons with BDNF (Leal, Comprido et al. 2017); (ii) stimulation of cultured hippocampal neurons and hippocampal synaptoneurosomes with BDNF induces the local synthesis of Pyk2 (Schratt, Nigh et al. 2004, Afonso, De Luca et al. 2019); (iii) downregulation of Pyk2 with a specific shRNA, or transfection with a dominant negative form of the kinase, abrogated the effects of BDNF on the synaptic expression of GluN2A in cultured hippocampal neurons (present study); (iv) transfection of hippocampal neurons with the wild-type Pyk2 mimicked the effects of BDNF and even synergized with the BDNF-TrkB signaling to further enhance GluN2A synaptic surface expression. It is interesting to note that the effects of BDNF on the surface expression of GluN2A- and GluN2B-NMDAR are both dependent on local translation and on Pyk2 but exhibit very distinct kinetics. This difference suggests that, in addition to shared mediators, other yet-unidentified mechanisms contribute differentially to the synaptic recruitment of the two receptor populations following TrkB receptor activation. A previous study showed that GluN2A, but not GluN2B, is dendritically translated and inserted in the postsynaptic membrane upon synaptic activity in the hippocampus (Swanger, He et al. 2013). Such a mechanism may also contribute, at least in part, to the translational activity required for BDNF to increase synaptic GluN2A content.

The results obtained in the experiments performed in cultured hippocampal neurons transfected with the dominant negative form of Pyk2 also showed that the kinase mediates the effects of BDNF on the extrasynaptic distribution of GluN2A and was required for the maintenance of this population of receptors on the dendritic plasma membrane under resting conditions. This suggests that the effects of BDNF on the synaptic expression of GluN2B-containing NMDAR may be secondary, at least in part, to the increase in their total surface expression.

PKC acts as a key mediator of BDNF-induced GluN2A surface expression, as Gö 6983 abolished the neurotrophin’s effect on synaptic GluN2A accumulation. TrkB receptors are coupled to activation of PKC via a phospholipase Cγ–dependent pathway (Leal, Comprido et al. 2014), and PKC has been shown to phosphorylate Pyk2 at Tyr402 (De Luca, Mele et al. 2025), a critical event for downstream signaling activation (Das, Pal et al. 2021, de Pins, Mendes et al. 2021). Therefore, these findings support a model in which BDNF-TrkB activation stimulates PKC, which in turn activates Pyk2 through phosphorylation. Dimerization and activation of Pyk2 is coupled to the stimulation of the tyrosine kinase Src (de Pins, Mendes et al. 2021), and GluN2A contains several tyrosine residues that can be phosphorylated by this kinase (Yang and Leonard 2001, Tahiri, Corti et al. 2025). The effect of BDNF in the up-regulation of surface GluN2A following their phosphorylation on specific tyrosine residues is thought to result from the inhibition in their interaction with the µ2 subunit of the AP2 protein adaptor complex, with the consequent association with endocytic clathrin-coated vesicles (Roche, Standley et al. 2001, Salter and Kalia 2004). Importantly, GluN2A Y1325 phosphorylation by Src was coupled to the facilitation of LTP induction in CA1 synapses (Taniguchi, Nakazawa et al. 2009, Yang, Trepanier et al. 2012). Although GluN2A subunits of NMDAR are also PKC substrates (Tahiri, Corti et al. 2025), this direct pathway is unlikely to mediate the BDNF-induced upregulation in GluN2A-NMDAR surface expression, as our results show that the effect is abolished by either Pyk2 downregulation or expression of a dominant-negative form of the kinase.

Our results also show that TrkB signaling contributes to the upregulation of GluN2A expression in a model of status epilepticus. In rats subjected to pilocarpine-induced seizures, GluN2A surface expression was increased in hippocampal synaptoneurosomes, an effect that was blocked by ANA-12, a selective TrkB inhibitor. Consistent with these observations, previous reports showed an increase in tyrosine phosphorylation of GluN2A after seizure activity in kainate-(Moussa, Ikeda-Douglas et al. 2001) and Li^+^/pilocarpine-induced status epilepticus (Niimura, Moussa et al. 2005). The increase in GluN2A phosphorylation in the latter model was correlated with an upregulation in activity of Src (Niimura, Moussa et al. 2005). Furthermore, studies performed in an in vitro model of epileptiform activity, displaying spontaneous epileptiform discharges in the CA3 region of rat hippocampal slices, showed an upregulation in Src activity associated with the epileptiform activity and a reduction in the frequency of epileptiform discharges upon pharmacological blockade of Src-family kinases (Sanna, Berton et al. 2000). The microelectrode array recordings performed in the present study further support the functional relevance of this signaling pathway in epileptogenesis. The application of a cocktail of drugs to induce epileptiform synchronization in cultured hippocampal neurons (bicuculline, 4-AP, and glycine) (Avoli, de Curtis et al. 2016) significantly increased burst frequency and network burst frequency, an effect that was impaired by either scavenging endogenous BDNF with TrkB-Fc or selectively inhibiting GluN2A with TCN 213. These results show a key role for GluN2A-NMDAR and BDNF in the activity-induced synchronization of hippocampal neuronal networks.

The observed upregulation in synaptic GluN2A-NMDAR associated with epileptogenesis is remarkable considering that inhibition of these receptors with NVP-AAM077 inhibited epileptogenesis in experiments using rats exposed to the kindling model or using the pilocarpine-induced epilepsy. In contrast, inhibition of GluN2B-NMDAR with ifenprodil was without effect in the development of seizures in both models (Chen, He et al. 2007). In the same study, GluN2A-NMDAR were preferentially coupled to the induction of the expression of the BDNF mRNA (Chen, He et al. 2007). Given the effects of BDNF to enhance the synaptic expression of GluN2A-NMDAR, this may constitute a positive feedback loop to enhance neuronal excitability.

Collectively, our findings define a novel BDNF-TrkB-dependent signaling mechanism that regulates the synaptic incorporation of GluN2A-containing NMDARs via the PKC-Pyk2-hnRNPK axis, requiring protein synthesis and dynamically modulated across different temporal windows and neuronal compartments. This pathway not only underscores the complexity of neurotrophin-mediated synaptic modulation but also provides new molecular targets for disorders characterized by dysregulated excitatory signaling, such as epilepsy. Future studies should investigate whether the molecular machinery identified here is similarly involved in activity-dependent plasticity phenomena such as long-term potentiation and whether pharmacological modulation of this pathway can yield functional benefits in animal models of synaptopathies.

## Acknowledgements

This work was financed by the European Regional Development Fund (ERDF), through the Centro 2020 Regional Operational Programme and the COMPETE 2020 Operational Programme for Competitiveness and Internationalization and Portuguese national funds via *Fundação para a Ciência e a Tecnologia* (FCT), under projects UIDB/04539/2020, UIDP/04539/2020 and LA/P/0058/2020. PDL was supported by FCT (PD/BD/135498/2018).

